# The nucleating agent crystallophore induces instant protein crystallization

**DOI:** 10.1101/2024.05.13.593864

**Authors:** Claude Sauter, Dominique Housset, Julien Orlans, Raphaël de Wijn, Kévin Rollet, Samuel Rose, Shibom Basu, Philippe Bénas, Javier Perez, Daniele de Sanctis, Olivier Maury, Eric Girard

## Abstract

The rapid preparation of homogeneous suspensions of micro- or nano-crystals is a crucial step in serial crystallography. We show how additives, such as the crystallophore (TbXo4) that acts as a molecular glue by promoting protein-protein interactions, can facilitate sample preparation for both serial synchrotron crystallography (SSX) and micro electron diffraction (3D ED). This lanthanide complex was used here for its nucleating properties to crystallize hen egg white lysozyme. SAXS monitoring indicates that crystals formed in a few minutes in low salt conditions that would not lead to spontaneous nucleation. Resulting micro- and nano-crystals were successfully used to determine the structure of the lysozyme-TbXo4 complex by SSX and 3D ED, illustrating the diffraction quality of the produced crystals and the usefulness of such compounds in the sample preparation pipeline for serial crystallography.

## INTRODUCTION

Obtaining well-diffracting crystals is a prerequisite to any diffraction-based experiments. Among the latter, macromolecular crystallography (MX) has provided valuable information at atomic scale on the architecture of biological molecules such as active site description, bound ligand interactions or details of interactions in key interfaces. Conventional MX relies on the use of a single crystal to obtain a complete crystallographic data set. However, the technique has been limited by radiation damages and most of the MX diffraction data collected during the past three decades were performed at cryogenics temperature to limit such damages. ^1,2^ It is worth noting that the averaged “static” structures provided by conventional MX may not be sufficient to fully understand the function of a given macromolecule in particular the molecular motions associated with the biological mechanism, such as catalytic mechanisms and associated conformational changes. To observe the macromolecules in motion at all the biologically relevant time scales, i.e. from the fs to a few minutes, we have to measure the diffraction data at room-temperature and find an alternative way to deal with radiation damages.

The application of serial crystallography (SX), at X-ray free electron laser facilites (XFEL) or at synchrotrons, provides an elegant way to overcome conventional MX limitations in particular for time-resolved (TR) experiments. Indeed, SX is based on the possibility to reconstruct a complete data set from single diffraction frames, each frame being measured from a single crystal. The inherent nature of SX data collection drastically lessens the radiation damage issue as it distributes the required X-ray dose over a very large number of crystals and opens the way to room or physiological temperature data collection, eliminating the tedious optimization of cryogenic conditions.

As a result, TR-SX allows to obtain structural dynamics information and observe biological macromolecules in action by capturing transient intermediates along a biological pathway. ^3–6^

From an experimental point of view, serial crystallography brings new constraints on crystal preparation as it intrinsically requires a large amount of samples to make sure to collect a complete diffraction data set. Moreover optimal time-resolved experiments require crystalline samples with a narrow size distribution in order to ensure a uniform triggering of the reaction under study through the entire crystal volume. ^7–9^

The ability to generate crystals on-demand may open new opportunities in TR-SX by facilitating rapid co-crystallization with substrates. First, it would enable the preparation of fresh crystals just prior to data collection, hence eliminating the issues associated with sample transportation, such as the control of the temperature, vibrations or mechanical shocks, or light for light-sensitive crystals. It would also reduce the loss of diffraction quality due to crystal ageing Second, it may also limit common soaking issues such as bad substrate/fragment diffusion due to crystal packing or interferences with crystal contacts. Indeed, molecules used for fragment based drug discovery may be incubated with the protein prior to crystallization, as can the substrates or analogues be prepared with the crystallization solution thus ensuring a controlled triggering or inhibition of the targeted reaction.

Recently, Henkel and co-workers proposed a method that combines protein crystallization and data collection in a novel one-step process they named JINXED to perform optimal time-resolved experiments based on serial crystallography. ^10^ They exploited hen eggwhite lysozyme (HEWL) crystals generated with crystallization times of only a few seconds from highly concentrated protein and crystallant solutions.

In 2017, we introduced the crystallophore (TbXo4), a lanthanide complex with both nucleating and phasing properties. ^11^ In particular, its unique characteristics of favoring protein crystallization was initially highlighted using a set of 8 proteins. The study also showed that the crystallophore induces more crystallization conditions for a given protein and strongly enlarges its crystallization diagram, allowing to use less concentrated protein solution. ^11^ These nucleating properties were also exploited to grow crystals from protein fractions containing a mixture of several proteins. ^12^ Finally, the addition of crystallophore to HEWL solubilized in pure water, instantly produced larger size particles that were detected by Dynamic Light Scattering (DLS). After several weeks, the presence of single crystals was observed in this unusual condition. ^13^

Altogether, these results prompted us to challenge the nucleating properties of the crystallophore for the minute-scale production of crystals with the appropriate size for either SX experiments or electron diffraction of 3D nanocrystals (3D ED). As a proof-of-principle, we generated HEWL crystals with a distribution ranging from the nanometer to the micrometer size allowing their diffraction quality to be evaluated by means of electron diffraction and synchrotron serial crystallography, respectively. Finally, we used time-resolved small-angle X-ray scattering (TR-SAXS)to determine the time window associated with the crystal production assisted by the crystallophore.

## MATERIALS AND METHODS

### Chemicals

The crystallophore molecule, TbXo4, was kindly provided by Polyvalan (https://crystallophore.fr). Hen egg white lysozyme was purchased from Roche (Cat. N° 10837059001) or from Seikagaku Corp. (Tokyo, Japan, Cat. N° 100940) or Sigma-Aldrich (St. Louis, MO, USA, Fluka Cat. N° 62970-5G-F). All other chemicals were purchased from Sigma-Aldrich.

### Sample preparation for diffraction experiments

The lyophilized hen egg-white lysozyme (Roche) was first dissolved in 80 mM sodium acetate pH 4.6 to a final concentration of 80 mg/mL. In a 1:1 ratio, the prepared solution was then mixed to a crystallant solution containing 800 mM NaCl, 80 mM sodium acetate pH 4.6 and 20 mM TbXo4 leading to a final volume of 100 μl. This batch crystallization results in a milky suspension. In the present work, crystallant refers to precipitating agent according to ^14^.

### Sample preparation for SAXS experiments

HEWL powders from Seikagaku Corp. or Sigma-Aldrich were dissolved in 50 mM Na acetate pH 4.5 to prepare stock solutions at 200 and 175 mg/mL, respectively. Lysozyme stock solutions were filtered prior to concentration measurement by UV spectrophotometry and ultracentrifuged at 125,000 x g prior to SAXS experiments.

### SSX chip preparation, data collection and data processing

After centrifugation during 2 minutes at 2000g, 5 μL of HEWL-Xo4 microcrystals supernatant solution was sealed between two foils of 13 μm thick Mylar (DuPont) enclosed in a frame of 10×15 mm (SOS chip). ^15^

Measurements were performed at the ESRF synchrotron facility (Grenoble, France) on ID29 beamline (https://www.esrf.fr/id29) using an X-ray photon energy of 11.56 keV (1.072 Å) with a 1% bandwidth and a beam size of 4 x 2 μm^2^ (H x V). X-ray pulse length of 90 μs at a frame rate of 231.25 Hz were recorded on a Jungfrau 4M detector. The sample-to-detector distance was 150 mm. The motorized stage was moved of 25 μm between two X-ray pulses to refresh the sample.

Diffraction patterns were processed using CrystFEL-v0.10.2 suite. ^16^ Spot-finding was performed using Peakfinder8 algorithm. ^17^ Various indexing algorithms, Xgandalf ^18^ and mosflm ^19^, were applied to index diffraction frames. Merging was done using partialator. ^20^ The final data set statistics are presented in Table 1.

**Table 1:** Data processing and refinement statistics for SSX data collected on HEWL microcrystals.

| Data processing |  |
| --- | --- |
| No. of collected frames | 153,600 |
| No. of hits | 40,262 |
| Indexed patterns | 22,622 |
| Average Dose per pulse (MGy) | 0.66 |
| Wavelength ($\text{\AA}$) | 1.072 |
| Space group | P4<sub>3</sub>2<sub>1</sub>2 |
| Unit-cell parameters ($\text{\AA}$) | a=b= 78.75, c= 38.20 |
| Resolution range (all. $\text{\AA}$) | 78.75 – 2.05 |
| High resolution range | (2.08 – 2.05) |
| No. observations | 2239287 (38381) |
| No. unique reflections | 7990 (380) |
| Multiplicity | 280.3 (101.0) |
| Completeness (%) | 100 (100) |
| R<sub>split</sub> | 11.87 (181.81) |
| CC<sub>1/2</sub> | 0.987 (0.213) |
| CC* | 0.997 (0.593) |
| SNR | 5.82 (0.50) |
| Wilson B factor ($\text{\AA}^2$) | 41.68 |
| Refinement |  |
| Resolution range | 55.68 - 2.05 (2.35 - 2.05) |
| Nb of refl. used in refinement | 7952 (2575) |
| Nb of refl. used for free set | 398 (129) |
| R<sub>work</sub> | 0.1801 (0.2807) |
| R<sub>free</sub> | 0.2166 (0.3018) |
| Number of non-hydrogen atoms | 1088 |
| macromolecules | 1001 |
| ligands | 30 |
| solvent | 57 |
| Average B-factor ($\text{\AA}^2$) | 44.29 |
| macromolecules | 43.39 |
| ligands | 68.22 |
| solvent | 47.64 |
| $\sigma$ bond ($\text{\AA}$) | 0.007 |
| $\sigma$ angle ($^\circ$) | 0.86 |
| Clashscore | 7.59 |

The HEWL structure was solved by molecular replacement with the PHASER program ^21^ from the CCP4 suite of programs, ^22^ using the 193L PDB entry as a starting model. The top 1 solution had an LLG value of 2274.4 and an R-factor of 0.361.

The model was further refined through two rounds of refinement in Phenix. ^23^ The first consisted in adding the Tb ion within the model using an anomalous Fourier synthesis and, in particular, by performing simulated annealing and water molecule search. Prior to the second refinement round, the Xo4 ligand was then added and the TbXo4 molecule occupancy was adjusted to 0.75 so that terbium B-factors fit ones of the binding residue Asp101. R-factors and stereochemistry values are summarized in Table 1.

### 3D ED grid preparation, data collection and data processing

The solution of lysozyme nanocrystals was centrifuged during 2 min at 2000 g and 3 μl of the supernatant were deposited on a 400 mesh copper grid covered with a Lacey carbon film. The grid was then backside blotted manually with blotting paper and flash frozen in liquid ethane using a home-made mechanical plunger.

The electron diffraction data were collected on a F20 cryo-electron microscope (FEI) operated at 200 kV and equipped with a hybrid pixel Medipix3RX 512×512 detector. Electron diffraction crystallographic data were collected at -170 °C on 5 different crystals (Table S1). A selected aperture of about 1 μm was inserted in the first image plane of the electron microscope in order to limit the area contributing to the measured diffraction pattern. The lysozyme crystals were continuously rotated on an angular wedge ranging from 32° to 46° with 0.1835° per frame (continuous rotation speed set to 5%) and an exposure time of 0.125 s/frame. The camera length was set to D= 730 mm, corresponding to calibrated camera length values of 1285 mm.

All the data sets were processed with the XDS package. ^24^ The 5 data sets were scaled and merged with XSCALE. The final data set statistics are presented in Table 2.

**Table 2:** Data processing and refinement statistics for electron diffraction data collected on HEWL nanocrystals.

| Data processing |  |
| --- | --- |
| Space group | P4<sub>3</sub>2<sub>1</sub>2 |
| Unit-cell parameters ($\text{\AA}$) | a=b=78.63 c=37.91 |
| Wavelength ($\text{\AA}$) | 0.02508 |
| Resolution range (all. $\text{\AA}$) | 55.60 – 3.21 |
| High resolution range | (3.30 – 3.21) |
| No. observations | 26621 (837) |
| No. unique | 1817 (123) |
| Multiplicity | 33.1 (11.8) |
| Completeness (%) | 83.8 (71.5) |
| $R_{\text{sym}}$ | 0.709 (1.941) |
| $R_{\text{meas}}$ | 0.736 (2.103) |
| $CC_{1/2}$ | 0.935 (0.393) |
| $I/\sigma(I)$ | 3.79 (0.69) |
| Wilson B factor ($\text{\AA}^2$) | 68.7 |
| Refinement |  |
| Resolution range | 50.00 – 3.21 (3.29 – 3.21) |
| No. of refl. (work set) | 1633 (102) |
| No. of refl. (free set) | 178 (9) |
| Number of atoms | 1033 |
| R-factor | 0.203 |
| $R_{\text{work}}$ | 0.189 (0.291) |
| $R_{\text{free}}$ | 0.331 (0.448) |
| $\sigma$ bond ($\text{\AA}$) | 0.007 |
| $\sigma$ angle ($^\circ$) | 1.810 |

The hen egg white lysozyme structure was solved by molecular replacement with the PHASER program ^21^ from the CCP4 suite of programs, ^22^ using the 193L PDB entry as a starting model. The top 1 solution had an LLG value of 540 and an R-factor of 0.454. The solution was further refined with Refmac5, ^25^ using electron atomic scattering length tabulated in the atomsf_electron.lib file of CCP4, leading to an Rwork of 0.202 and an Rfree of 0.318, with good stereochemistry (r.m.s. bond and angle deviation of 0.06 Å and 1.84°, respectively). At this point, the first peak in the residual {*F*_*obs*_ *– F*_*calc*_} Coulomb potential map (+7.32 σ) was observed at the expected position of the TbXo4 compound (Figure 2A). TbXo4 was thus added to the model and placed in the position and orientation that provided the best fit with the residual Coulomb potential map. Another round of refinement with Refmac5 indicated that the TbXo4 B-factor was significantly higher than those of Trp62, Trp63 and Asp101 side chains, that are in close contact with TbXo4. The occupancy of TbXo4 was then reduced to 0.75 and a final round of refinement was performed. The final structure included the entire polypeptide chain corresponding to the hen egg white lysozyme and one TbXo4 compound, with acceptable R-factors and stereochemistry (Table 2).

### SAXS sample environment and data collection

TR-SAXS experiments were performed on the SWING beamline at synchrotron SOLEIL (Saint-Aubin, France). The X-ray beam wavelength was set to λ = 0.826 Å (15 keV). The Eiger X 4M detector (Dectris) was positioned at a distance of 2,037 mm from the sample with the direct beam off-centered. The resulting exploitable q-range was 0.004 – 0.69 Å^-1^, where the wave vector is q = 4π sinθ / λ and 2θ is the scattering angle.

Rapid mixing of HEWL solutions with salt and nucleant solutions was performed using a home-made syringe pump as described in Figure 1. The first syringe (1 ml, Braun Melsungen AG, Germany) contained 50 mM sodium acetate pH 4.5 with or without HEWL, the second contained 50 mM sodium acetate pH 4.5, 400 or 800 mM NaCl, with or without 20 mM TbXo4. Injection lines made of Peek tubes coming from the syringes were coupled to the incubation capillary by a mixing tee (U-466S, Chrom Tech). The incubation capillary was directly plugged to the SAXS cell.

**Figure 1.**
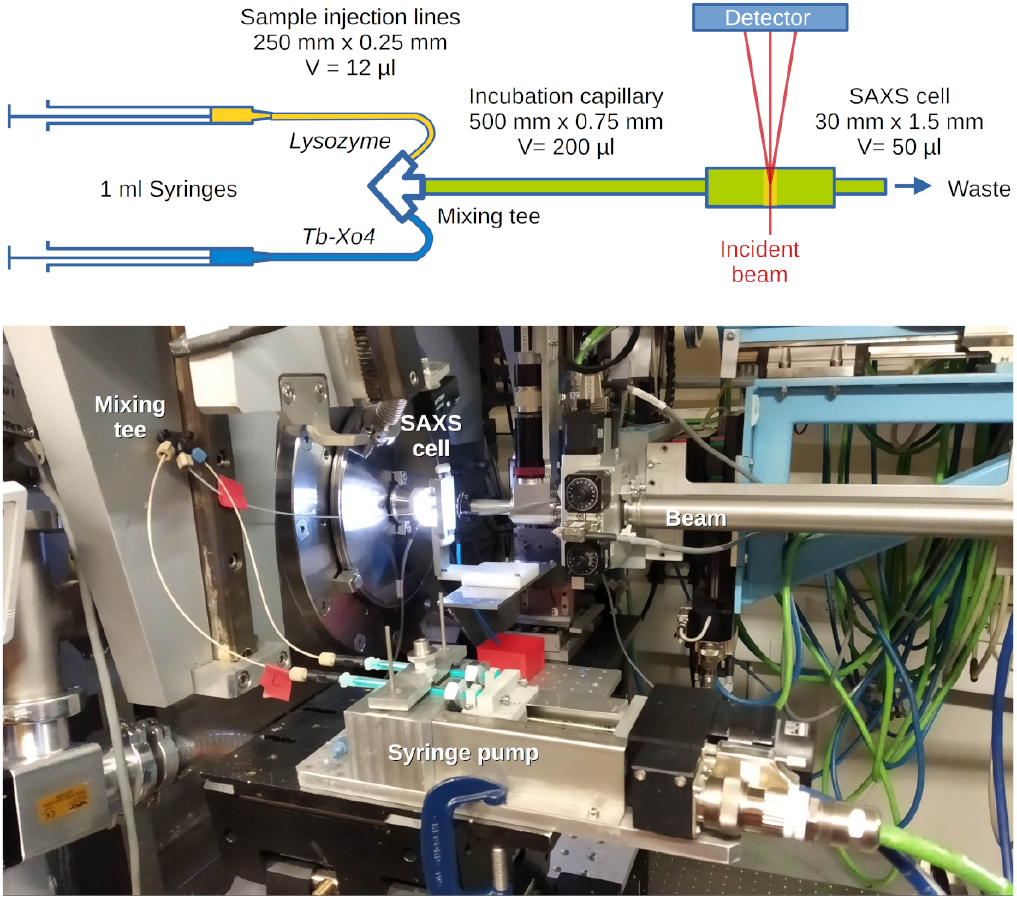
Schematic view and picture of the experimental setup used for TR-SAXS.

The syringes had an inner diameter of 4.73 mm, hence delivering 17.6 μl/mm by linear displacement of step motor. They were filled with 300 μl of solution before each experiment. Rapid solution mixing was carried out by moving the plunger of both syringes by 10.9 mm in 1 min 50 sec (0.1 mm/sec) to obtain 280 μl of crystallization medium and fill the entire tubing, including the SAXS cell. The evolution of the solution was monitored over 30 min by collecting 180 images every 10 sec, while translating the solution to refresh the sample (displacement step of 0.03 mm at 1 mm/sec) after each image (exposure 1 sec).

For each salt condition (final NaCl concentration 200 or 400 mM), a series of measurements was carried out including i) the acetate buffer only (for buffer subtraction), ii) control solutions containing either HEWL or Tb-Xo4, iii) the full mix to follow the nucleation/growth process. After each experiment the setup was extensively washed with a 2% (v/v) aqueous solution of Hellmanex (Hellma HmbH, Germany) and water.

Data processing was performed using Foxtrot ^26^ and data analysis with the ATSAS package. ^27^ Graphics were prepared with BioXTAS-RAW. ^28^

Powder diffraction programs were not suitable for Bragg rings indexing due to the large unit cell and the limited experimental resolution. Instead, Fcalc were generated for the experimental resolution range using Phenix. ^23^ Structure factors were calculated up to a resolution of 12.7 Å (2θ = 3.8°) from a HEWL crystal structure collected at room temperature (PDB ID 4NGK; a = 79.28 Å, c = 37.73 Å). Bragg rings overlayed on the SAXS scattering profile were assigned to their corresponding Bragg peaks by successive refinement of cell parameters by least-square minimization in an Excel spreadsheet converged to a = 79.0 Å and c = 38.3 Å.

### Raw data availability

The raw frames for both SSX and 3D ED experiments have been deposited in Zenodo repository under DOI name DOI1 and DOI2 for SSX and 3D ED data, respectively (*DOIs will be completed upon publication*).

Structural biology software support for IBS was provided by SBGrid._29_

## RESULTS AND DISCUSSION

### Rapid HEWL crystallization

The crystallization of HEWL, a widely used model protein for MX, in the presence of NaCl and acetate buffer at pH 4.5 is a conventional condition to generate well-diffracting crystals. In the present work, the mixing in an Eppendorf tube of a HEWL protein solution prepared in acetate buffer with a solution containing the crystallophore in the presence of NaCl (see Materials and methods section) yielded a dense milky suspension after less than 2 minutes. This milky suspension that slowly sediments is indicative of a potential massive presence of microcrystals. ^9^

### Crystal size distribution

Few microliters of the produced suspension were first deposited between two glass lamellas for optical visualization. As shown on Figure 2A, crystals with the typical tetragonal shape were clearly visible with a distribution size ranging from one micron to twenty microns. To characterize even smaller ones, Transmission Electron Microscopy (TEM) was performed and clearly revealed well defined crystals with size ranging from 200 nm to one micron (Figure 3A). This confirms the effect of TbXo4 on the production of crystalline samples with ideal size for both 3D ED and serial crystallography, providing optimal crystal size according to the planned experiments. It should be mentioned that the produced crystals were obtained at low NaCl concentration (that would not allow spontaneous crystallization) and commonly used HEWL concentration (40 mg/mL) thanks to the TbXo4 properties, and were obtained in batch facilitating the scaling up to larger volumes. As already mentioned, the crystallophore provides new crystallization conditions from nanolitre vapor diffusion crystallization. ^11^ By exploiting these conditions, one can expect to facilitate the sometime difficult transition from nano-scale crystallization to large-scale batch crystallization. ^7,8^

**Figure 2.**
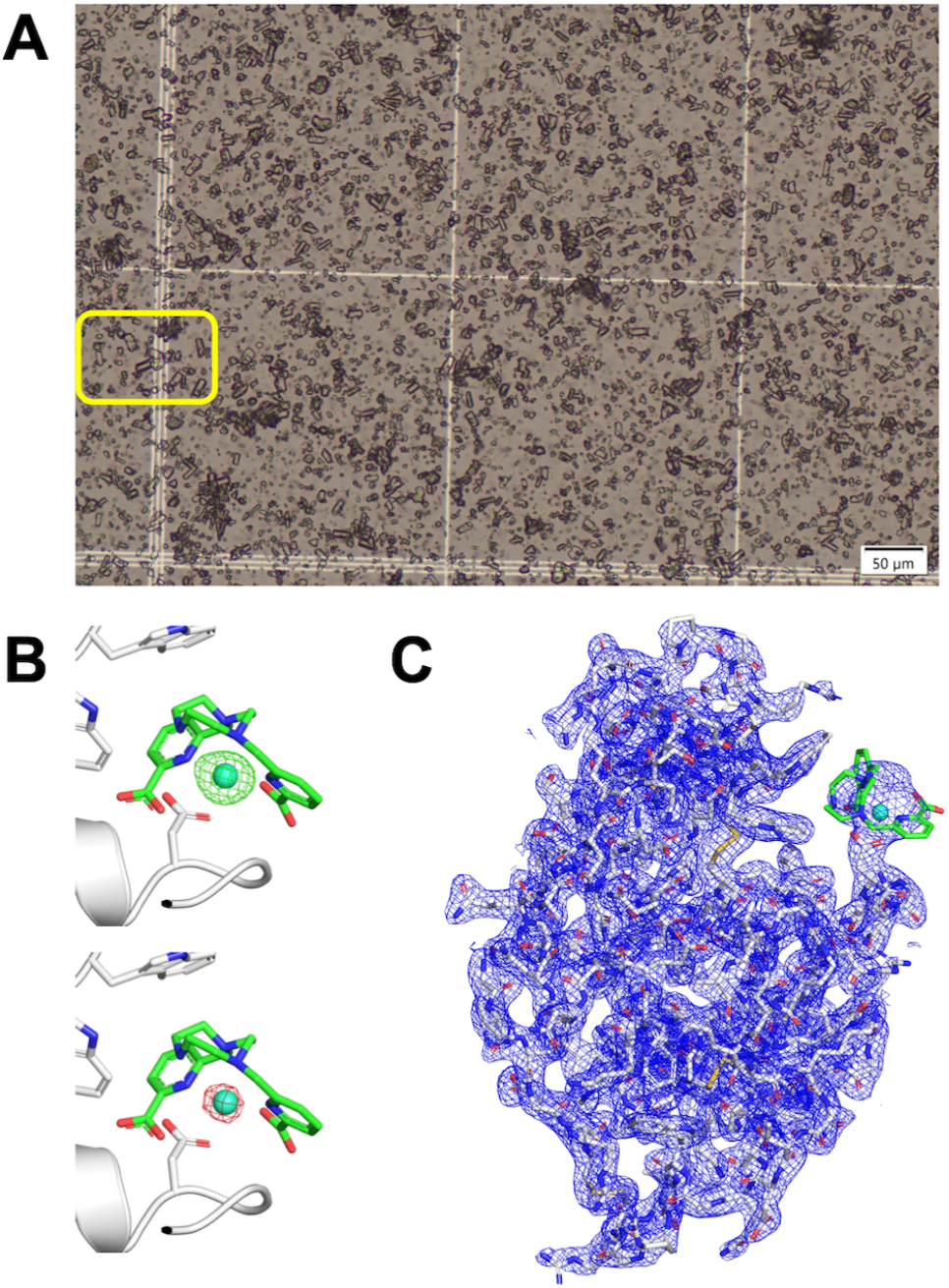
A) View of the HEWL micro-crystals used for the SSX experiments. A close-up (yellow rectangle) is provided in Supporting Information Figure S1. B) Electron density maps (contour: 7 σ level) calculated after molecular replacement, {*mF*_*obs*_*-DF*_*calc*_} in green (up) and anomalous Fourier synthesis in red (down). Refined HEWL (cartoon mode) model with TbXo4 depicted in green (stick mode) was superimposed. C) Refined HEWL model with {*2mF*_*obs*_*-DF*_*calc*_} electron density map (contour: 1 σ level).

**Figure 3.**
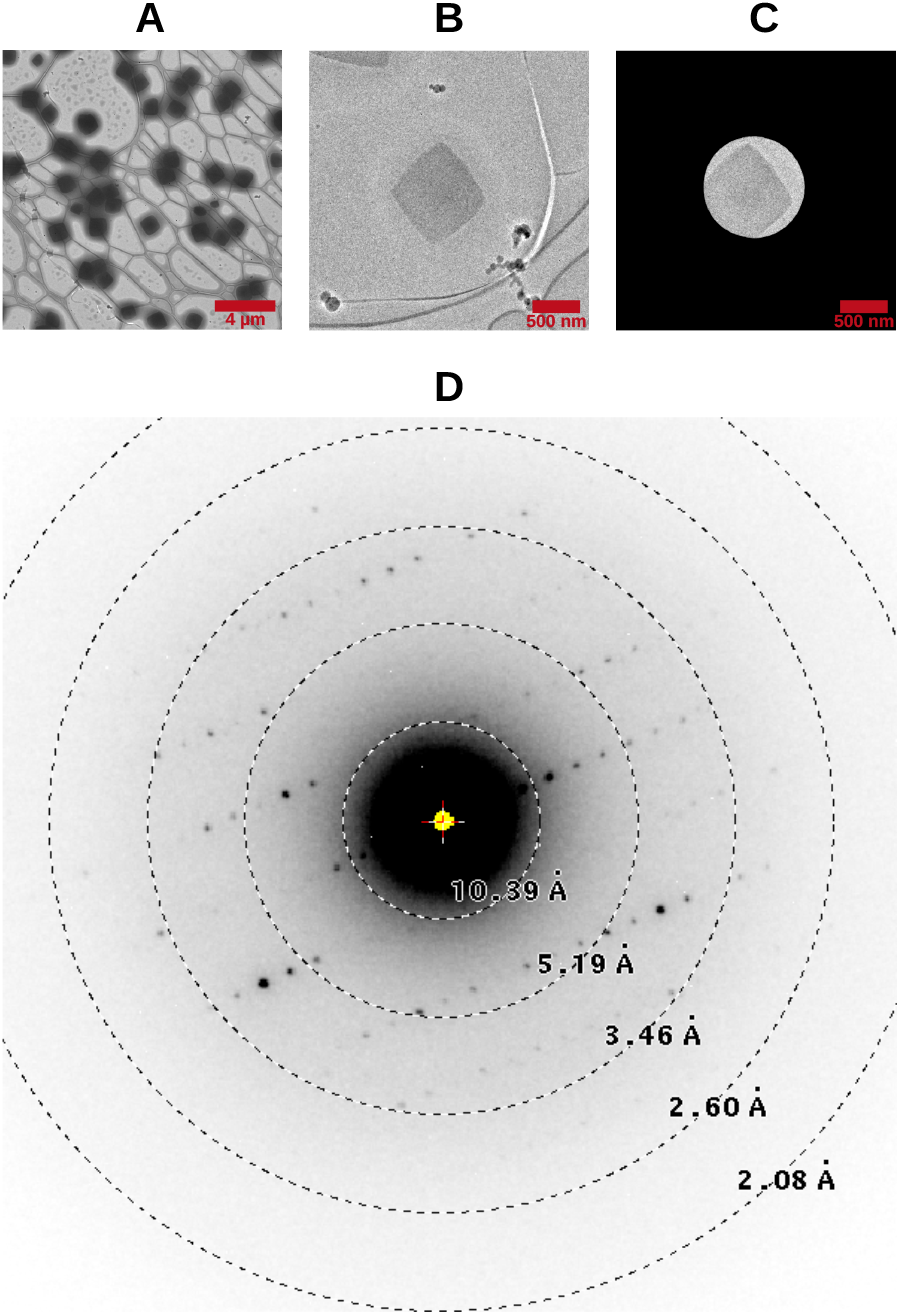
A) View of the HEWL crystals on the frozen grid that was used for the electron diffraction experiments. The magnification of the F20 electron microscope was set to x800. B) and C) View of the crystal that was used for the data collection of set no. 5. The magnification of the F20 electron microscope was set to x5000. View C) has been taken with the selected aperture diaphragm inserted. D) Still diffraction pattern of the crystal used for data set no.4, just before launching the continuous rotation.

### Diffraction evaluation of micron-sized crystals

A Synchrotron Serial Crystallography (SSX) experiment was performed on ID29 beamline at ESRF (Grenoble) using a fixed-target approach. To this end, a SOS (sheet-on-sheet) chip was used to present the crystals in front of the beam. ^15^ The solution containing micro-crystals was sandwiched between two Mylar foils (see Materials and Methods section) leading to random crystal positions. The diffraction data were recorded by defining a raster grid of 25 μm and by using a beam size of 4 x 2 μm^2^. 153600 detector frames were recorded. Details of data collection are described in the Material and Methods section. Among all recorded detector frames, 40,262 contained diffraction (hit rate of 26.2%) and 20622 of those contained indexable patterns (indexing rate of 51.2%).

The overall data processing resulted in a 2.1 Å resolution data set with 100% completeness, CC(1/2) and SNR of 0.987 and 5.82, respectively (Table 1).

As shown in Figure 2, the molecular replacement performed with Phaser yielded a clear {*2mF*_*obs*_*-DF*_*calc*_} electron density map (contour level XT = 1.0). In particular, in {*mF*_*obs*_*-DF*_*calc*_} electron density map, the presence of a 18 XT peak closed to Asp101 confirms the presence of the crystallophore molecule as expected. ^11^ This is furthermore confirmed by the anomalous Fourier synthesis exploiting the anomalous signal present within the data. Indeed, at the beam energy used to collect the data (11.56 keV), the terbium f” value is about 8 electrons thus inducing a significant anomalous contribution to the diffracted intensities (Figure 2B).

These results clearly highlight the quality of the TbXo4 generated micro-crystals and their usefulness to derive structural information at high resolution.

### Diffraction evaluation of nano-sized crystals

3D ED is an emerging diffraction techniques allowing to get structural biology information from nano-sized protein crystals. ^30–32^ To evaluate the diffraction quality of the nano-sized HEWL crystals, we performed a 3D ED data collection using a F20 cryo-electron microscope from FEI (see Materials and Methods section). On the best grid, the crystal size distribution was moderately heterogenous, with a majority of crystals resembling square platelets, with edges ranging from 0.5 to 1 μm (Figure 3A).

The final merged dataset was obtained from 5 different crystals (Figure 3B and 3C). The individual data processing resulted in similar diffraction qualities for each sample with, in particular, high resolution comprised between 4 and 3.2 Å (Supporting Information Table S1). One example of diffraction frame is shown in Figure 3D.

The final data set is 83.8% complete and include data up to 3.21 Å resolution (Table 2).

After molecular replacement and the first round of refinement, the main peak in the residual {*F*_*obs*_*-F*_*calc*_} Coulomb potential map (+7.32 σ) was unambiguously attributed to the terbium atom (Figure 4A). As for the structure derived from SSX data, its position closed to Asp101 is as expected. ^11^ The final fit between the {*2F*_*obs*_*-F*_*calc*_} Coulomb potential map and the final model is good all along the polypeptide chain as well as for TbXo4 (Figure 4B) showing that TbXo4 generated nano-crystals allow to get precise structural information. It is worth to note that the occupancy determined for the TbXo4 moiety is the same for both the SSX and the 3D ED data derived model. This demonstrates the consistency of the data and the reproducibility of the crystals grown in this condition.

**Figure 4.**
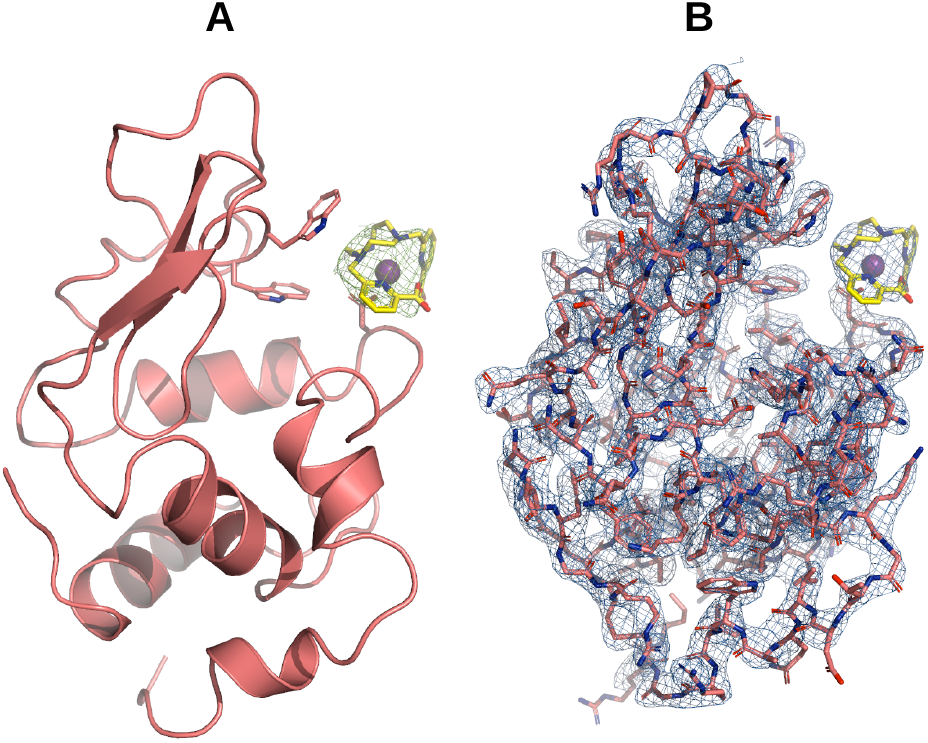
A) View of the HEWL structure after the first round of Refmac5 refinement during which the crystallophore TbXo4 was not included. The main peak of the residual {*F*_*obs*_ – *F*_*calc*_} Coulomb potential map contoured at the 2.8 σ level (depicted in green) is shown, with the model of TbXo4 shown in stick mode fitted into it. B) View of the {2*F*_*obs*_ – *F*_*calc*_} Coulomb potential map, contoured at the 1.1 σ level, after the final round of Refmac5 refinement including the TbXo4 compound.

### Crystal production time window assessed by SAXS

sTime-resolved SAXS was used to monitor the nucleation properties of TbXo4 on HEWL solutions. To perform rapid mixing of the crystallization solution with the HEWL, we assembled an injection system consisting of a syringe pump, a connector (tee-mixer) for the capillary tubes bringing the solutions from the syringes, and a wider capillary tube acting as an incubation chamber connected to the SAXS cell (Figure 1). Several procedures were tested to achieve rapid mixing without creating bubbles in the system. The best compromise consisted of moving the motor at 0.1 mm/s, resulting in a mixing step of 1’50” to produce a volume sufficient to fill the entire tubing, including the SAXS cell. The evolution of the solution was then followed for 30 min in taking one image every 10 sec and moving the solution after each image to refresh the sample. Also the beamline energy was shifted from its standard value (12 keV – λ = 1.033 Å) to 15 keV to avoid fast capillary fulling with solutions containing TbXo4.

Rapidly after mixing HEWL with 10 mM TbXo4 in 400 mM NaCl the solution turned snowy in the SAXS cell (Supporting Information Movie S1). The scattering intensity jumped at t = 170 s (Figure 5A) and discrete peaks appeared in the profiles, indicating the formation of crystalline material. The cell then gradually filled with microcrystals (Figure 5C), leading to powder diffraction patterns that intensified over time (Supporting Information Movie S2). Figure 5B illustrates the initial HEWL scattering signal and its superposition with microcrystal diffraction patterns after 25 min. The analysis of these powder diffraction-like patterns indicates the presence of tetragonal HEWL crystals with unit-cell parameters a = 79.0 Å and c = 38.3 Å (Supporting Information Figure S2).

**Figure 5.**
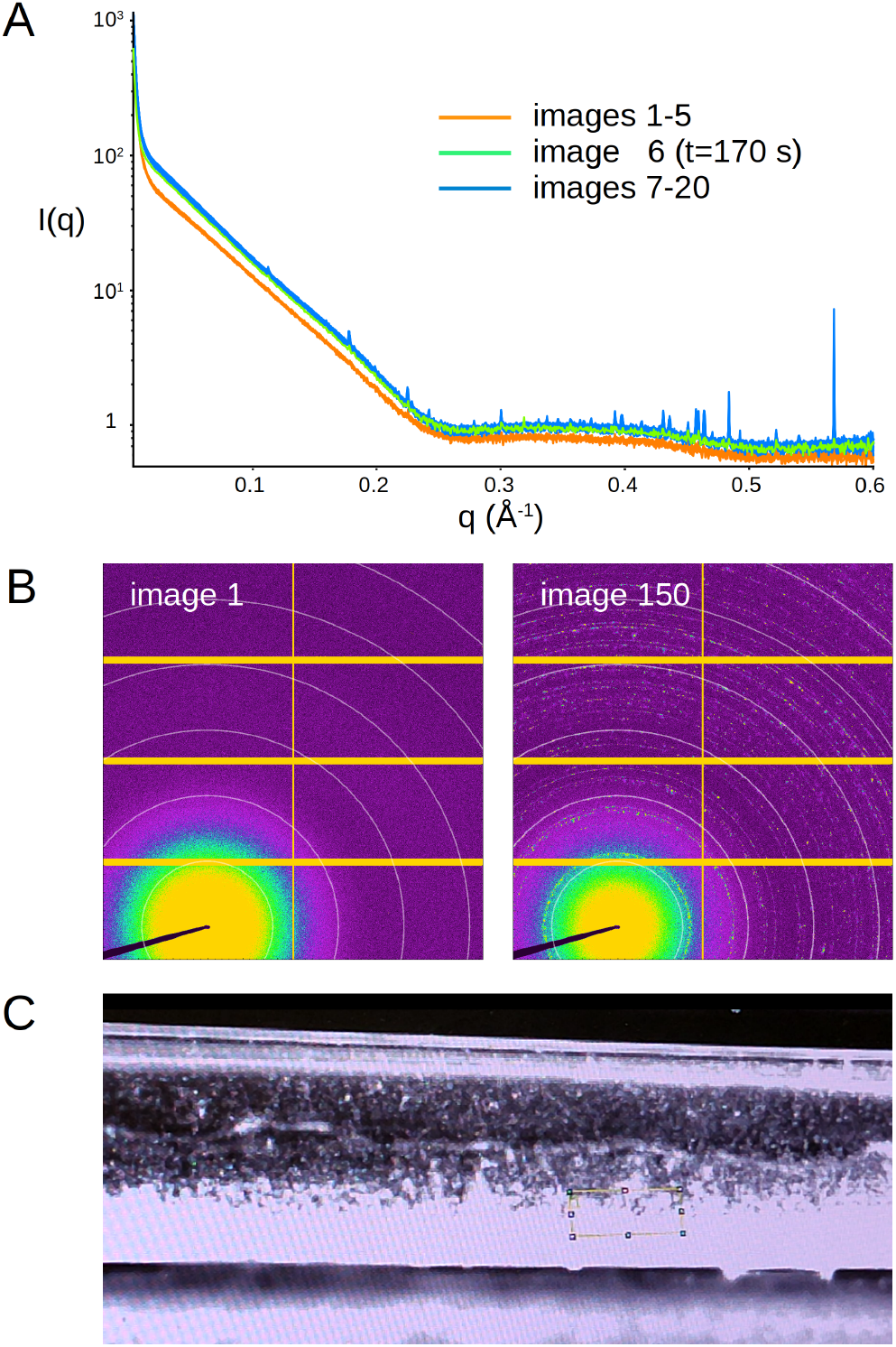
A) Monitoring of microcrystallization by TR-SAXS. A) Evolution of SAXS profiles between t=120 s (image 1) just after the mixing step and t=380 s (image 20). The scattering intensity increases and diffraction rings (Bragg rings) appear at t=170 s, indicating the formation of microcrystals. B) First image showing the scattering signal of HEWL solution, which superimposes with diffraction rings generated by microcrystals in image 150 (t= 1620 s). White circles correspond to a scattering angle of q = 0.1, 0.2, 0.3, 0.4, 0.5 and 0.6 Å^-1^, respectively. C) View from the beamline camera showing microcrystals settling to the bottom of the SAXS cell. The yellow rectangle symbolizes the X-ray beam.

Rapid crystallization did not occur in the time frame of the experiment (30 min) in the absence of TbXo4 or when lowering the NaCl concentration to 200 mM (data not shown). However, lowering the protein concentration to 67 mg/ml (instead of 100 mg/ml) still led to microcrystallization, although less spectacular (Supporting Information Movie S3).

## CONCLUSION

The production of protein crystals requires the generation of a supersatured state, which is usually achieved by adding a crystallant that reduces the solubility of the protein. In the case of lysozyme, the most common crystallant is NaCl. The higher the crystallant concentration, the higher the supersaturation and the higher the probability to nucleate and grow crystals. This is why batch crystallization with fast mixing of protein with high crystallant concentration is a very popular way of preparing sample for SSX which requires hundreds of thousands of microcrystals to determine a crystal structure. However, this kind of highly supersaturated conditions are very unstable because far from equilibrium and do not guarantee the reproducibility and the homogeneity of microcrystals batches.

Starting from the observation that TbXo4 could trigger the nucleation and the crystal growth of HEWL even in the absence of any crystallant, ^13^ we explored the possibility to generate micro-crystal batches at low salt concentrations, not by making the protein less soluble but by promoting intermolecular contacts leading to crystal growth. This is exactly what TbXo4 provides, as illustrated by its propensity to generate more hits at lower protein concentration during crystallization screening. ^11^

Our results illustrate how TbXo4 can initiate fast microcrystal production in a few minutes. SAXS monitoring reveals the presence of diffraction patterns after less than 3 min. However, owing to the large beam size and the strong background from the solution, we cannot exclude that nanocrystals may appear much faster but remained undetectable until they reach a minimal size and density in the SAXS cell. Nevertheless, such microcrystalline samples are easily and reproducibly prepared on demand. They proved useful for structural analysis by SSX and 3D ED, and showed a diffraction quality that is fairly homogeneous.

Altogether, when we consider the possibility to promote instant crystallization with TbXo4, along with its ability to trigger crystallization in different solvent conditions of a screen and generate multiple crystal forms, the crystallophore definitely appears as a promising tool for sample preparation in SSX including time resolved applications.

## ASSOCIATED CONTENT

### Supporting Information

The Supporting Information is available free of charge on the ACS Publications website.

**Table S1:** Lysozyme data acquisition by 3D ED and integration statistics of individual nanocrystals (PDF file).

**Movie S1**: Evolution of HEWL – TbXo4 mix over time. The first sequence was recorded after 3 min when microcrystals started to appear and sediment at the bottom of the SAXS capillary. The solu-tion was pushed forward every 10 s by the syringe pump to refresh the sample exposed to the X-ray beam symbolized by the yellow rectangle. The second sequence at 12 min after the mix shows increased micro-crystal formation and sedimentation (mp4 video format).

**Movie S2**: Sequence of 150 SAXS images (1 image every 10 s) illustrating the appearance and intensification of diffraction rings with a mix consisting of 100 mg/ml HEWL, 10 mM TbXo4, 400 mM NaCl, 50 mM Na acetate pH 4.5 (mp4 video format).

**Movie S3**: Sequence of 180 SAXS images (1 image every 10 s) illustrating the appearance and intensification of diffraction rings with a mix consisting of 75 mg/ml HEWL, 10 mM TbXo4, 400 mM NaCl, 50 mM Na acetate pH 4.5 (mp4 video format).

## AUTHOR INFORMATION

### Author Contributions

CS and EG designed and coordinated the project. / RdW, KR, PB, OM and EG prepared the samples. DH and EG performed 3D ED experiments / DH processed and analyzed 3D ED data / JO, SR, SB, DdS and EG performed SSX data collection / JO and EG processed and analyzed SSX data / CS, KR, PB, JP and EG performed SAXS data collection/ CS and PB processed and analyzed SAXS data / CS, DH and EG wrote the original draft / The final manuscript was written through contributions of all authors. / All authors have given approval to the final version of the manuscript.

### Notes

OM and EG declare a potential conflict of interest since they are cofounders of the Polyvalan company that commercializes the crystallophore.

## ACKNOWLEDGMENT

IBS acknowledges integration into the Interdisciplinary Research Institute of Grenoble (IRIG, CEA). The authors thank the following synchrotron facilities and associated scientists for beamtime allocation to the project: beamline ID29 at the European Synchrotron Facility (Grenoble, France), SWING at SOLEIL synchrotron (project 20211662, Saint-Aubin, France). We thank Emmanuelle Neumann for training on the F20 cryo-electron microscope.

## ABBREVIATIONS

MX: macromolecular crystallography –
HEWL: hen egg-white lysozyme –
SSX: synchrotron serial crystallography – 3D
ED: electron diffraction of 3D nanocrystals.

## SUPPORTING INFORMATION

**Table S1:** Lysozyme data acquisition by 3D ED and integration statistics of individual nanocrystals.

| Data set | 1 | 2 | 3 | 4 | 5 |
| --- | --- | --- | --- | --- | --- |
| Wavelength (Å) | 0.02508 |  |  |  |  |
| Exposure (s/frame) | 0.125 |  |  |  |  |
| $\Delta\phi$ (°/frame) | 0.1835 | | | | |
| Number of frames | 206 | 253 | 208 | 252 | 175 |
| $\Phi_{\text{total}}$ (°) | 37.8 | 46.4 | 38.2 | 46.2 | 32.1 |
| Camera length (mm) | 1285 | 1285 | 1285 | 1285 | 1285 |
| Data processing |  |  |  |  |  |
| Space group | **P4<sub>3</sub>2<sub>1</sub>2** |  |  |  |  |
| Unit cell dimensions | a=b=78.5<br>c=38.9 | a=b=78.5<br>c=38.8 | a=b=78.3<br>c=38.7 | a=b=78.8<br>c=37.0 | a=b=78.5<br>c=38.9 |
| Resolution (Å) | 35.0 – 3.32<br>(3.52 – 3.32) | 27.8 – 3.21<br>(3.40 – 3.21) | 55.4 – 3.41<br>(3.62 – 3.41) | 24.9 – 3.21<br>(3.33 – 3.21) | 78.5 – 4.01<br>(4.25 – 4.01) |
| Reflections | 5521 (870) | 7546 (1172) | 5048 (767) | 5834 (379) | 2650 (415) |
| Unique reflections | 1394 (219) | 1566 (338) | 1132 (173) | 1427 (109) | 667 (104) |
| Completeness (%) | 69.5 (68.9) | 71.0 (70.7) | 61.6 (61.3) | 67.2 (49.8) | 57.1 (56.8) |
| R<sub>sym</sub> | 0.518 (1.991) | 0.564 (1.505) | 0.698 (1.992) | 0.451 (1.385) | 0.713 (0.995) |
| R<sub>meas</sub> | 0.596 (2.275) | 0.635 (1.687) | 0.792 (2.256) | 0.513 (1.626) | 0.826 (1.151) |
| I/σI | 2.54 (0.88) | 2.44 (0.77) | 2.04 (0.62) | 3.38 (0.84) | 1.55 (1.09) |
| CC<sub>1/2</sub> | 0.891 (0.425) | 0.905 (0.408) | 0.909 (0.293) | 0.921 (0.438) | 0.733 (0.550) |

**Figure S1.**
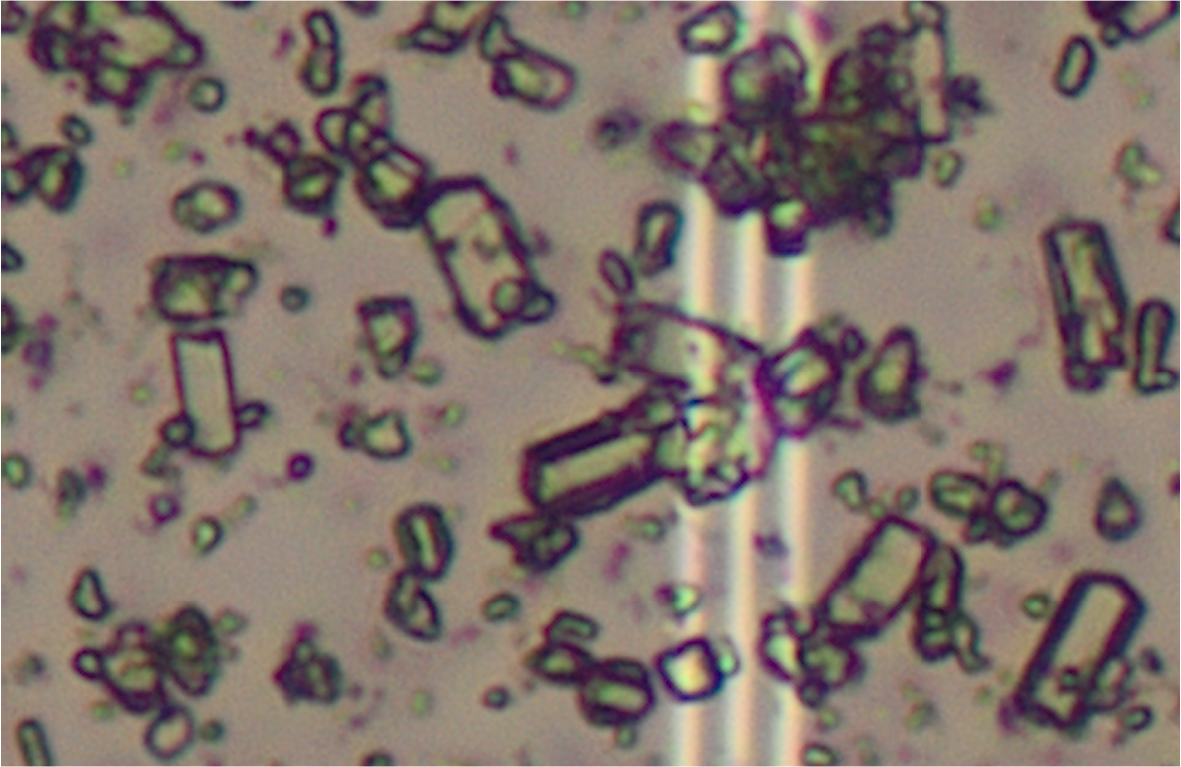
Close up on HEWL microcrystals used for the SSX experiments (PDF file).

**Figure S2:**
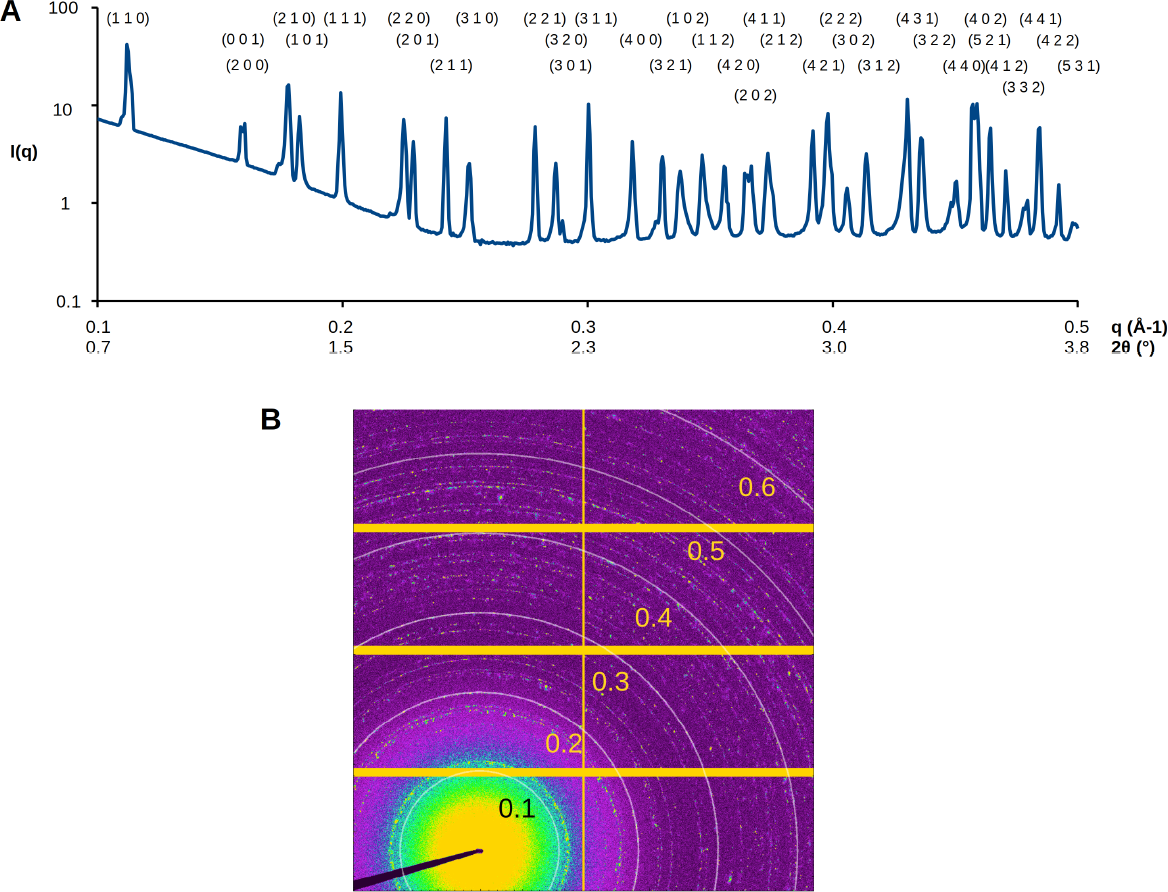
A) SAXS profile averages over images 135 to 150 highlighting powder diffraction ring positions as a function of q and 2θ. Miller indices of peaks are indicated and correspond to tetragonal microcrystals with cell parameters of a = b = 79.0 Å and c = 38.3 Å. B) Image 150 showing the superposition of diffuse scattering from the solution and powder diffraction signals from microcrystals. q-values corresponding to resolution rings are indicated (PDF file).

